# Joint and Distinct Cognitive–Motor Associations with Multivariate White Matter Microstructure and Connectivity

**DOI:** 10.1101/2024.06.05.597645

**Authors:** Zaki Alasmar, Stefanie A. Tremblay, Thomas R. Baumeister, Felix Carbonell, Claudine J. Gauthier, Yasser Iturria-Medina, Christopher J. Steele

## Abstract

Recent advances in cognitive neuroscience emphasise the importance of healthy white matter (WM) for optimal behavioural functioning. It is now widely accepted that brain connectivity via WM contributes to the emergence of behaviour. However, the association between the microstructure of WM fibres and behaviour is poorly understood due to the indirect and overlapping nature of methods used to assess microstructure. Here, we used the Mahalanobis Distance (D2) to integrate 10 metrics of WM derived from multimodal neuroimaging that have strong ties to microstructure. The D2 metric was chosen because it accounts for metrics’ covariance as it measures the voxelwise distance between every subject and the average; thus providing a robust multiparametric assessment of microstructure. We used multivariate correlation to examine voxelwise WM-behaviour associations with two cognitive and two motor tasks, which allowed us to compare within and across behavioural domains. We observed that behaviour is organised in cognitive, motor, and integrative components that have widespread associations with WM, from frontal to parietal regions. Notably, the decomposition of these factors shows that tasks traditionally labelled as cognitive also contain motor components that map onto motor-related WM patterns, and that motor tasks likewise contain non-motor components linked to distinct WM microstructural features. Our results highlight the complex nature of the links between microstructure and behaviour, and support the relevance of multivariate modelling when examining brain-behaviour associations.

## 1. Introduction

Recent advances in neuroimaging techniques allow for the non-invasive assessment of brain structure and function in health and disease <u>(Calhoun, 2018; Tardif et al., 2016)</u>. These techniques have become crucial in understanding the neural underpinnings of cognitive and motor behaviour. Though most research has focused on brain function, and thus functional imaging, in relation to behavioural tasks, it is now increasingly clear that function is supported by the connectivity and microstructure of white matter (WM) <u>(Thiebaut de Schotten et al., 2020;</u> <u>Thiebaut de Schotten & Forkel, 2022)</u>. WM can be assessed with diffusion weighted magnetic resonance imaging (DWI), providing both an estimation of WM connectivity architecture and a quantification of different components of its microstructure <u>(Duval et al., 2017; Soares et al.,</u> <u>2013)</u>. Importantly, there are many quantitative DWI-derived metrics that are differentially sensitive to various aspects of microstructure (e.g. organisation, myelination, and density) <u>(Lv et</u> <u>al., 2025)</u>, thereby serving as a rich multivariate source of individual microstructural differences that can be directly related to behaviour. Investigating how WM microstructure is related to normative behaviour is crucial for enhancing our basic understanding of brain function and a first step in investigating how pathologies affect structure and behaviour and how they may be ameliorated by targeted treatment <u>(Thiebaut de Schotten & Forkel, 2022)</u>.

WM fibres vary in their diameter, myelination, density, and overall composition <u>(Mezer</u> <u>et al., 2013; Stikov et al., 2015)</u>. Changes in healthy WM are largely due to refinements in the signal conduction environment <u>(Hagmann et al., 2010)</u> to support optimal function <u>(Pajevic et</u> <u>al., 2014)</u>, resulting in observable behavioural effects such as enhanced/reduced performance. Myelination is known to shape this environment by fine tuning signal conduction <u>(Knowles et</u> <u>al., 2022)</u>, with animal models providing key evidence for myelin remodulation during motor learning and cognition (e.g. in memory; <u>McKenzie et al., 2014; Pan et al., 2020)</u>. While there is a large body of work in human neuroimaging focused on cortical function and structure, there has been much less work to explore how the microstructure of the underlying WM connections support behaviour. Expanding from initial cross-sectional human work (e.g., <u>Bengtsson et al.,</u> <u>2005; Johansen-Berg et al., 2007</u>), neuroplastic changes in DWI-derived measures of WM have been shown to be associated with a wide range of behaviours, including motor skills <u>(Scholz et</u> <u>al., 2009; Steele et al., 2013; Taubert et al., 2010; Tremblay et al., 2021)</u>, memory (Alasmar et al., 2026; de Lange et al., 2017; Johansen-Berg et al., 2007), and development <u>(Muetzel et al.,</u> <u>2008, 2018)</u>. Thus, individual differences in WM microstructure that arise as a function of individual experience can be assessed with non-invasive MRI and used to probe the relationship with behaviour in both healthy and disease states. Previous research has largely relied on assessing individual contrasts with bivariate approaches (most commonly assessing fractional anisotropy [FA] and mean diffusivity [MD]), which assumes independence when DWI metrics cannot be directly tied to a single microstructural or physiological source <u>(Carter et al., 2025;</u> <u>Raghavan et al., 2021; Tardif et al., 2016)</u>. In contrast, in multivariate frameworks, each metric can be considered as an indirect measure that provides complementary information for a more comprehensive characterization of WM microstructure (Parent et al., 2025) . In such cases, the cumulative insight from different metrics provides a more sensitive marker of the underlying microstructure (Tremblay et al., 2024).

DWI-derived metrics provide measurements of WM microstructure such as tissue integrity and fibre characteristics (Raffelt et al., 2017; Raghavan et al., 2021; Tardif et al., 2016). Simple and complex diffusion models can be computed from DWI multi-shell high angular resolution protocols <u>(Jelescu & Budde, 2017)</u>, such as the diffusion tensor (DTI) <u>(Soares et al.,</u> <u>2013)</u>, fibre orientation distribution function (FOD; <u>Tournier et al., 2019)</u>), and neurite orientation dispersion and density imaging (NODDI; <u>(Zhang et al., 2012)</u>); which can then be used to assess microstructural properties such as myelin integrity, fibre thickness, and intracellular volume. While the metrics derived from DTI, FOD, and NODDI have been unquestionably useful in human neuroscience and often interpreted as specific physiological parameters <u>(Jones et al., 2013)</u>, there is no clear one-to-one mapping between metrics and physiological parameters. Further, we and others have shown that there is a high degree of correlation between metrics within and across models (Carter et al., 2025; Figley et al., 2022; Tremblay et al., 2024; Tremblay et al., 2025; Tremblay et al., 2025; Uddin et al., 2019), suggesting that strong conclusions based on theoretical interpretation of a single metric are often difficult to justify.

Human behaviour is often measured with task-specific assessments that are typically interpreted as reflecting a single cognitive construct or domain. However, this overlooks the multiple interacting brain systems that give rise to behaviour (Varoquaux & Poldrack, 2019), and how behavioral tasks imperfectly sample these domains (e.g., reading and motor tasks both engage visual pathways). And while standard univariate brain–behaviour analyses ignore this shared covariance (Poldrack, 2010; Varoquaux et al., 2018), multivariate approaches can be used to reveal latent factors (Schöttner et al., 2023) that are more directly tied to brain structure and function (Price & Friston, 2005; Varoquaux et al., 2018). The advantages of multivariate approaches have been demonstrated in recent work in memory and motor behaviour (Pur et al., 2022), as well as in cognition (Voigt et al., 2023; Ziegler et al., 2013). Similar issues arise in studies of WM where most analyses rely on individual microstructural metrics that potentially have multiple physiological determinants <u>(Liao et al., 2024; Tardif et al., 2016)</u>, limiting interpretability of tissue-specific contributions. To address this, we recently developed an integrative multimodal framework for quantifying individual differences in WM microstructure by combining highly correlated, physiologically related, metrics while accounting for their covariance (utilizing the Mahalanobis distance: D2; Tremblay et al., 2024). Previous work has used D2 to identify WM abnormalities in traumatic brain injury (Taylor et al., 2020), relate epilepsy duration to WM tract alterations (Owen et al., 2021), and, more recently, by our group to characterize individual variation in the corpus callosum (Tremblay et al., 2024) and relate WM microstructure to cardiovascular health (Tremblay et al., 2025; Tremblay et al., 2025).

In the present study, we aimed to identify the unique and overlapping spatial patterns of the relationship between WM microstructure and cognitive and motor behaviour. We used the Mahalanobis distance (D2) to integrate MRI metrics while accounting for their shared covariance and performed a partial least squares (PLS-SVD) multivariate statistical analysis to relate multivariate behaviour from four cognitive and motor tasks to voxel-wise white matter D2. We hypothesised that integrating multiple measures of microstructure and behaviour together yields more biologically interpretable and meaningful brain-behaviour associations, and that there would be differential patterns in D2’s relationship to behaviour across WM.

## 2. Method

### a. Participants

All data was sourced from the WU-Minn Human Connectome Project (HCP S1200 release). The details for the HCP data acquisition and preprocessing are described in depth elsewhere (Van Essen et al., 2013). Briefly, our sample consisted of the minimally preprocessed data from 1065 healthy young adults (age: M= 28.76 years, SD= 3.68; 556 females), with no history of psychiatric, neurological, or neuropsychological disorders, and no history of substance abuse. We excluded participants with incomplete imaging data and/or invalid acquisitions. The HCP acquired informed consent prior to any data collection. All data used in this study was published with consent to publish from the HCP.

### b. Multimodal Neuroimaging Protocols

All imaging was conducted by the HCP on a custom 3T Connectome Skyra MRI scanner with a 32-channel head coil. Anatomical scans were acquired in the first session, and included T1-weighted as well as T2-weighted protocols (Van Essen et al., 2013). The T1w scans were acquired using a 3D-MPRAGE sequence while a 3D T2-SPACE sequence was used for T2w. Anatomical scans were acquired with a 0.7mm isotropic resolution (FOV=224x224). T1w had a TI=1000, TE=2.14, and TR=2400, while T2w TE was 565 and TR of 3200. Mult-shell DWI data (TE/TR=89.5/5520 ms, FOV=210×180 mm, b-values of 1000, 2000 and 3000 s/mm^2^ and a 1.25 mm isotropic resolution, 90 uniformly distributed directions, and six b=0 volumes) were collected in the fourth session. Full acquisition details can be found at: https://www.humanconnectome.org/hcp-protocols-ya-3t-imaging. Sixty-four participants were excluded in the current study due to poor cerebellar coverage, resulting in a final sample of 1001 subjects.

### c. Microstructural features from multimodal neuroimaging

#### Neuroimaging preprocessing

All MRI scans were preprocessed following the minimal preprocessing procedure by the HCP (Glasser et al., 2013), which included intensity normalisation, and eddy current and susceptibility-induced distortions corrections using the reverse phase encoding of b=0 scans. This was followed by co-registration of DWI to native T1w (rigid body) and b-vector reorientation, and motion and gradient nonlinearity correction with the HCP pipeline. All subsequent preprocessing steps and metric extraction are described in detail in Tremblay and colleagues (2024) and described briefly below.

First, the preprocessed multishell DWI was bias-field corrected using the ANTs’ N4 algorithm via dwibiascorrect from MRtrix3 (Tustison et al., 2010). Using MRtrix3’s dwi2tensor (Tournier et al., 2019), we calculated the diffusion tensor and extracted its metrics with tensor2metric, which yielded voxelwise fractional anisotropy (FA), mean diffusivity (MD), axial diffusivity (AD), and radial diffusivity (RD). DTI metrics maps were transformed into the group template space as described below. We then used the multi-tissue Constrained Spherical Deconvolution (CSD) method to estimate a WM fibre response function, and later WM fibre composition. We first segmented all tissue types (WM, GM, CSF) by applying MRtrix3’s 5ttgen (using the FSL algorithm, (Smith et al., 2012; Tournier et al., 2019) on T1w scans. We computed the response function of each tissue type for all participants from the minimally preprocessed DWI data (without bias field correction) and the five-tissue-type (5tt) image using the msmt_5tt algorithm of the dwi2response function (Dhollander et al., 2016, 2018; Jeurissen et al., 2014; Raffelt et al., 2017). The response function represents the diffusion profile of a specific fibre population and is used to estimate the Fibre Orientation Distribution Functions (FOD) with CSD. The WM, GM and CSF response functions were then averaged across all participants, resulting in a single response function for each of the three tissue types. Multi-shell multi-tissue CSD was then performed in each individual with the average response functions to obtain an estimation of FODs for each tissue type. This step is performed using the dwi2fod msmt_csd function of MRtrix3 within a brain mask. Bias field correction and global intensity normalisation, which normalises signal amplitudes to make subjects’ data comparable, were then performed on the FODs using the mtnormalise function in MRtrix3.

#### Group space coregistration

While most studies focus on optimising GM alignment, our WM focus necessitated a WM-specific registration. For this, we used a multi-contrast registration that was primarily driven by WM FODs but also included information about GM and CSF. We created population templates for WM, GM, and CSF FODs based on a subset of 200 participants using MRtrix3’s population_template function (nl_update_smooth= 1.0, nl_disp_smooth= 0.75). We subsequently nonlinearly registered all subjects and our population template using MRtrix3’s mrregister function using the same parameters, and then applied the computed warps to WM FODs, DTI metrics (FA, MD, AD, and RD), and brain masks with mrtransform. During this step, the WM FODs were transformed without reorientation, which is described below. To create a template mask encompassing only the voxels with data from all subjects, we computed the intersection of all warped brain masks. Additionally, we warped the WM probability images from the five-tissue-type (5tt) segmentation to the group template space to generate a group WM probability mask. Lastly, we used the T1w and T2w images in their native resolution (0.7 mm^3^ isotropic) to calculate the T1w/T2w ratio to generate a proxy myelin map, which was then warped to the FOD template (Glasser & Van Essen, 2011).

The FOD template was then segmented to extract the WM fibre elements (fixel) mask. Fixel segmentation was then performed on the WM FODs of each subject and the fixels were normalized and aligned to the template using the subject-to-template warps (fixelreorient) and mapped to the corresponding fixels in the fixel mask (fixelcorrespondence function). This ensured a consistent set of fixel directions for all subjects. We computed 3 fixel metrics: apparent fibre density (FD), fibre bundle (FC: expansion/contraction relative to the template), and fibre density and cross-section (FDC: FD*FC, overall capacity of a fibre bundle to carry information). The fixelwise measures were converted to voxelwise maps for inclusion in the multimodal model by computing FD as the sum of the FOD lobe integrals, FC as the FD-weighted mean of FC using the fixel2voxel mean option so that higher-density fibre populations contributed proportionally more to the aggregate value, and FDC as the voxelwise sum of FDC using the fixel2voxel sum option to yield a cumulative estimate of microstructural capacity within each voxel.

The bias field-corrected DWI data was also fit with the neurite orientation dispersion and density imaging (NODDI) model using the python implementation of Accelerated Microstructure Imaging via Convex Optimization (AMICO) including all b-values (0, 1000, 2000, 3000) (Daducci et al., 2015; Zhang et al., 2012). The response functions were computed for all compartments, and the fitting procedure was performed on the unbiased DWI volumes within the brain mask. The resulting parameters obtained from the fitting process were the intracellular volume fraction (ICVF) assessing neurite density (i.e. interpreted as axonal density in WM) and orientation dispersion index (OD) assessing the extent of dispersion around the mean orientation.

In total, 10 voxelwise microstructural measures were computed for each of the 1001 subjects to use as features in our analyses. These features were: 4 DTI measures (FA, MD, RD, AD), 3 FOD measures (FC, FD, FDC), 2 NODDI measures (ICVF & OD), and T1w/T2w.

### d. Multivariate distance model

We used the Mahalanobis Distance (D2), a voxelwise multivariate distance implemented within MVComp (https://github.com/neuralabc/mvcomp) (Tremblay et al., 2024), to generate individual subject D2 maps quantifying the distance of each WM voxel from the reference group mean. We applied a leave-one-out approach to avoid biasing D2 by removing each subject from the reference population used to compute their comparison mean and covariance. The output maps (D2) represent, for each subject, a voxelwise assessment of the multivariate differences in WM microstructure between that subject and the reference population (i.e. all other subjects).

### e. Outlier detection

We removed every participant with extensive voxelwise outliers, based on a threshold of 5 standard deviations from their average D2 value. That is, if a participant’s D2 map contained 55 or more voxels with D2 values larger than 5 SD (i.e., > 0.07% voxels) of their own voxelwise average, that participant was dropped from the analysis. This value was chosen as the optimal tradeoff between outlier removal and retaining a large sample as further increasing the %-voxels cutoff results in a ∼50% drop in sample size. As D2 is computed relative to the average, we reran the D2 computations on the resulting final smaller sample of 735 subjects. We then applied a power transformation at each voxel (across all subjects) to normalise the D2 distribution and z-scored it for standardisation. Lastly, we winsorised the resulting matrices of standardised D2 z-scores with probability threshold of 99.7% (∼3 SD) in each tail.

### f. Behavioural tasks

We used 4 tasks to assess cognition and motor functioning from the NIH toolbox included with the HCP (Reuben et al., 2013; Weintraub et al., 2013). To assess motor functioning, we used the Grip Strength task (GST) and the 9-Hole Pegboard task (9-HPT), examining strength and dexterity respectively. In the GST, subjects were asked to squeeze a hand dynamometer as hard as they could, while in the 9-HPT, they were timed while placing and then removing 9 pegs in holes on a board as quickly as possible. The tasks chosen to assess cognition were the card sorting task (CST) and the list sorting task (LST), both well validated and standardised tasks widely used to assess cognitive flexibility and working memory respectively. All behavioural measures were also power-transformed and z-scored. The voxelwise correlations between these tasks and D2 are provided for completeness in Figure 1.

### g. Age and sex correction

To account for the potentially confounding effects of age and sex (Weber et al., 2022), each of the behavioural data, as well as the voxelwise D2 values, were fitted with a robust linear regression (task ∼ age + sex, D2 ∼ age + sex) and the residuals retained for subsequent analysis to statistically remove their effects. As with the imaging data, age and sex measures were power-transformed and z-scored prior to inclusion in the analyses.

### h. Statistical analyses

A multivariate PLS-SVD (MATLAB version R2022a; MathWorks) was used to relate voxelwise D2 to the four behavioural measures (GST, 9-HPT, CST, LST) by decomposing the brain–behaviour covariance matrix into latent variables that maximize shared variance (Lin et al., 2022). For a covariance matrix M, SVD yields M=USV^T^, where columns of U give voxel weights and rows of V^T^ give task weights for each latent variable. Statistical significance of latent variables was tested with 1000 permutations by shuffling behavioural data and comparing the explained covariance to the empirical solution. Voxel and task weight reliability was assessed with 1000 bootstrap resamples, rerunning the decomposition each time to derive confidence intervals. Significance was set at family-wise error–corrected p<0.05.

Significant findings were further assessed to identify their WM structural connectivity profiles using a normative model of whole-brain connectivity, given the impact of connectivity on cortical areas (Alasmar et al., 2025). For each significant LV, we chose a maximum of 3 regions of spatially connected voxels with large weights in order to individually identify the set of streamlines passing through them. This approach is identical to that used in identifying the network effects of brain lesions (Kaeja et al., 2026; Karnath et al., 2018; Talozzi et al., 2023; Thiebaut de Schotten et al., 2020; Zayed et al., 2020), and provides a visualization of the connectivity for each of the identified white matter regions. Here, we used a previously constructed streamline connectivity model that consisted of 10 million representative streamlines between all GM regions in our template space (Kaeja et al., 2026; Parent et al., 2025). After extracting the streamlines passing through the clusters, the GM connectivity profile of the WM passing through these clusters was summarised using the Automated anatomical labelling atlas 3 (AAL; Rolls et al., 2020) with the tck2connectome function of MRTrix3. A schematic of the statistical procedure is illustrated in Figure 1.

## 3. Results

We used D2 and PLS-SVD to examine the multivariate relationship between microstructure and cognitive and motor behaviour. We found that the first 3 latent variables (LVs) were statistically significant, explaining 39.01%, 25.23%, and 19.04% of brain-behaviour covariance, respectively (Figure 1). The PLS scores of D2 and behaviour were positively correlated in all LVs (r=0.39, r=0.69, r=0.70 for LV1, LV2 and LV3, respectively; panel A, Figures 2, 3, 4). LV1 was characterised by positive behavioural weights on both cognitive tasks (cognitive flexibility and working memory) and one of the two motor tasks (manual dexterity) (Figure 2B). Significant voxels were distributed in WM underlying frontal and parietal regions and the cerebellum, and were characterized by predominantly negative weights, indicating that the lower D2 (closer to the reference group) was associated with better performance (Figure 2C). LV2 exhibited strong positive weights with grip strength and weaker positive weights with cognitive flexibility, and was primarily characterized by WM voxels with positive and negative values in both parietal and cerebellar WM (Figure 3B,C). Lastly, LV3 showed strong positive weights with manual dexterity and weaker negative weights for working memory (Figure 4B). LV3 was characterized by predominantly positive weights in WM (frontal and corpus callosum), with additional strong negative weights in frontal regions in close proximity. (Figure 4C). The detailed results for each LV are presented in the following sections.

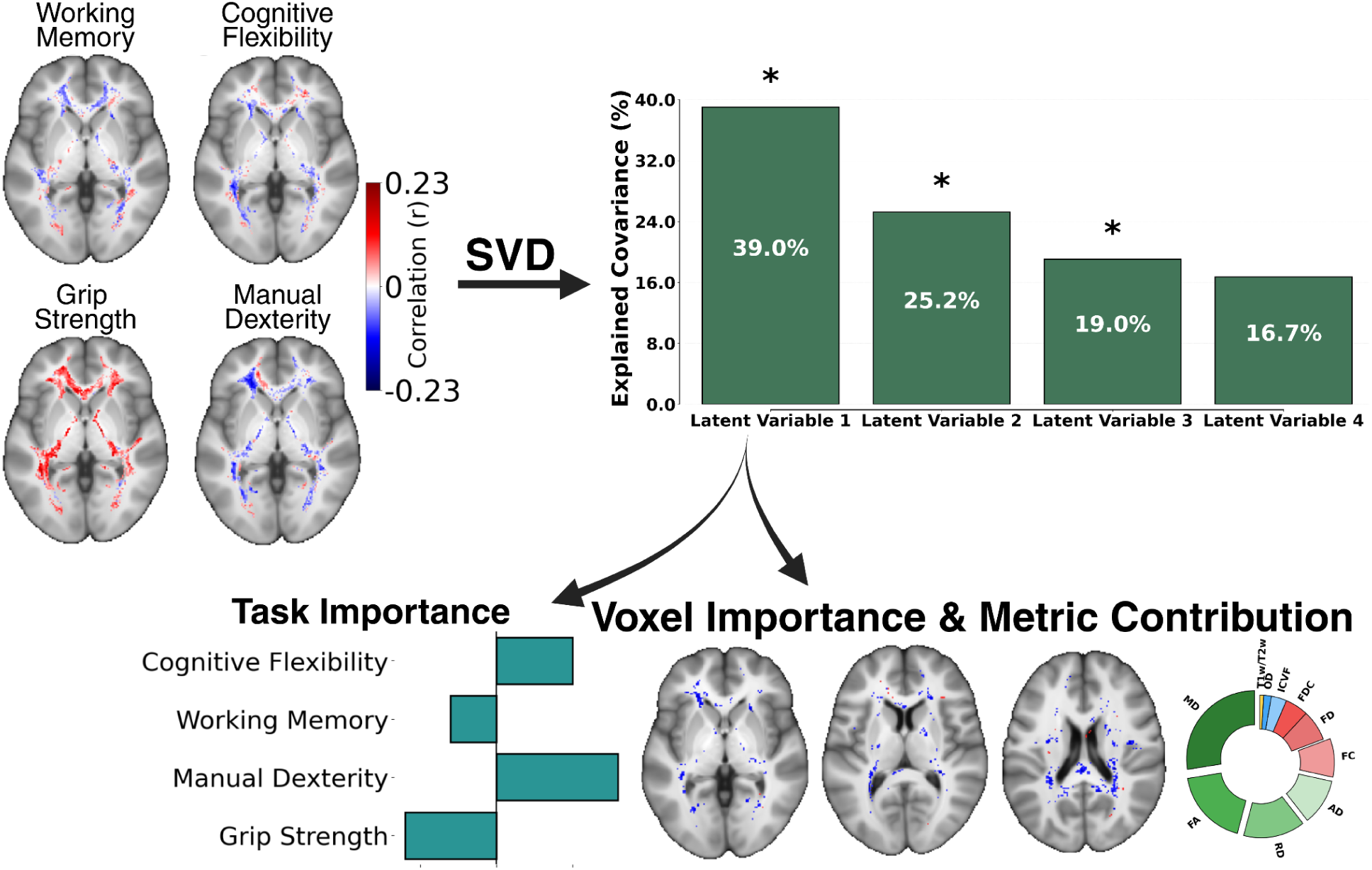

### a. Latent Variable 1

The first LV revealed a moderate association between transformed D2 and behavioural scores (r = 0.39, p < 0.001; Figure 2A). We found that cognitive flexibility, working memory, and manual dexterity were statistically significant in their positive weights (Figure 2B). WM voxels adjacent to frontal, temporal, and cerebellar regions had negative weights, while parietal regions had both negatively (notably, the corticospinal tract) and positively weighted regions (positive only in WM underlying the pericentral sulcal regions) (Figure 2C). Put differently, these distributed negative voxels across the WM along with the positively weighted tasks are negatively correlated given their opposite signs, while positively weighted voxels are positively correlated with behaviour.

Further probing identified three clusters with the largest spatially contiguous voxel weights, located in WM underlying the left pericentral sulcus, left prefrontal WM, and the right corticospinal tract. Across all three clusters, FC contributed most strongly to the clusters’ D2 scores (Figure 2D). We then examined the cortical connectivity profiles of these clusters to characterise the network-level basis of their respective WM–behaviour associations (Figure 2E). For the pericentral cluster, the greatest proportion of connections passed through the left supplementary motor area (SMA), followed by the right SMA. The prefrontal WM cluster showed strongest connectivity with the left putamen, followed by the left inferior frontal orbital cortex. Finally, streamlines passing through the corticospinal tract cluster were most strongly connected to the right thalamus, with additional prominent connections to the right parietal cortices and the right precuneus.

**Figure 2:**
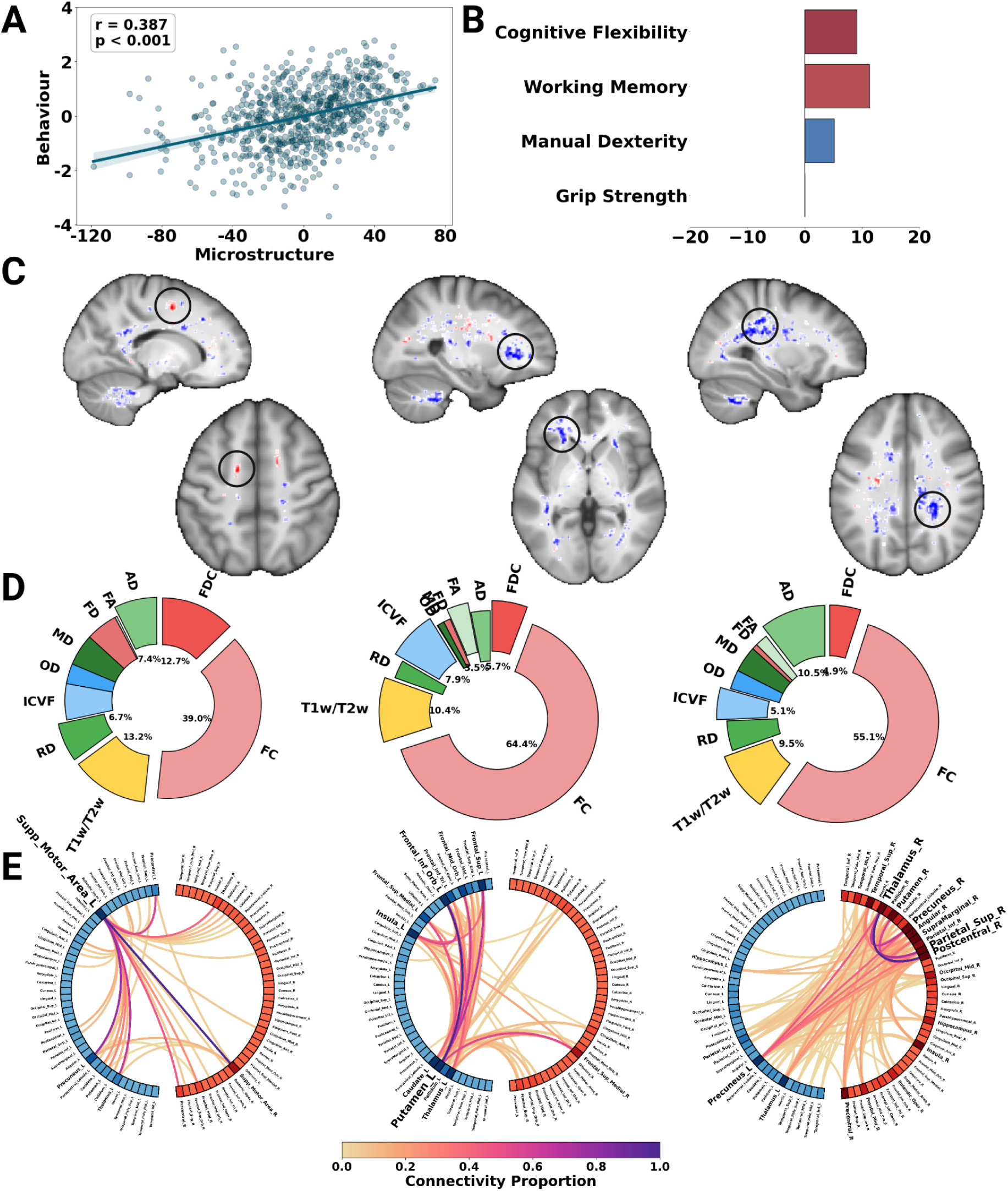
Decomposition results for the first latent variable. (A) PLS score correlation; (B) weights of statistically significant tasks; (C) weights of statistically significant voxels; (D) metric contributions to the D2 score; (E) connectivity profiles of spatially contiguous clusters based on the AAL atlas.

### b. Latent Variable 2

LV2 had a stronger association between PLS scores than LV1 (D2 and tasks; r = 0.69, p < 0.001; Figure 3A). Both grip strength (largest weight) and cognitive flexibility (smaller) were positively weighted, while manual dexterity had a small negative weight (Figure 3B). Unlike the more focal nature of LV1, there were positively- and negatively- weighted voxels distributed in WM (Figure 3C). The positively-weighted voxels were in the left parietal, right prefrontal, right temporal, and right superior cerebellar WM, whereas negatively weighted voxels were located in left frontal, callosal, and inferior cerebellar WM. The positive voxels in parietal and frontal regions are maximally positively correlated with cognitive flexibility and grip strength, but they are negatively correlated with manual dexterity. On the other hand, the negative voxels in frontal and colossal WM are negatively associated with cognitive flexibility and grip strength, but positively associated with manual dexterity.

We extracted three clusters located in the genu of the corpus callosum, left corticospinal tract, and WM underlying the right primary motor area. These clusters were characterised by dominant contributions from FC and T1w/T2w metrics (Figure 3D). When examining these clusters’ connectivity (Figure 3E), the genu of the corpus callosum emerged as a major hub of interhemispheric connectivity, primarily linking frontal cortices and the caudate nuclei. The connectivity profile of the corticospinal tract cluster was again dominated by left thalamic and parietal regions, and as expected, streamlines passing through the cluster underlying M1 predominantly influenced connectivity of M1 itself, followed by the right thalamus and the putamen.

**Figure 2:**
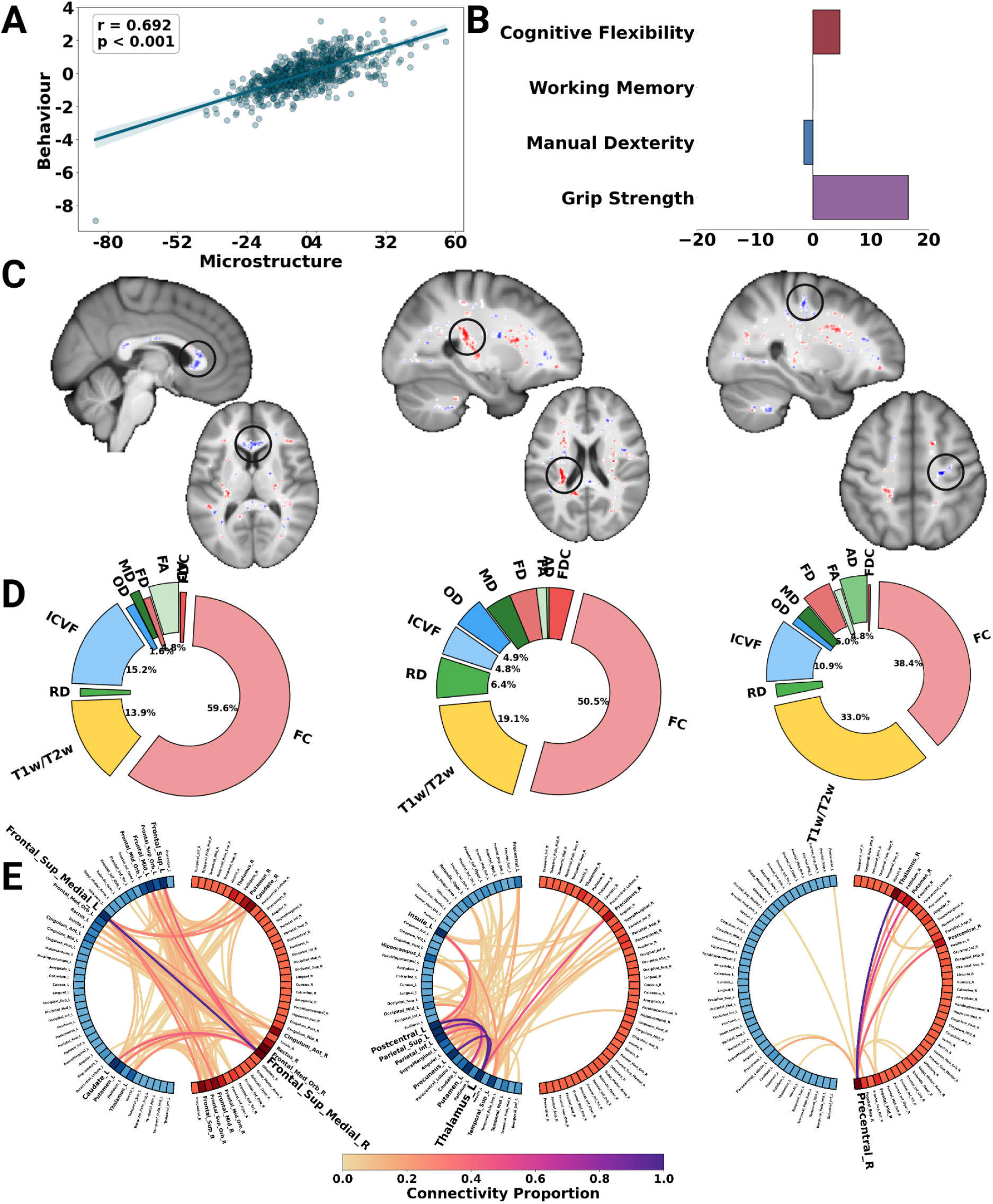
Decomposition results for the first latent variable. (A) PLS score correlation; (B) weights of statistically significant tasks; (C) weights of statistically significant voxels; (D) metric contributions to the D2 score; (E) connectivity profiles of spatially contiguous clusters based on the AAL atlas.

### c. Latent Variable 3

LV3 was also characterized by a strong correlation with behaviour (r = 0.70, p<0.001; Figure 4A). Working memory and manual dexterity weighed heavily on this LV, though in opposite directions (negative and positive, respectively) (Figure 4B). There were also negatively- and positively-weighted voxels distributed throughout WM. Significant positive voxels were mainly in prefrontal WM, and existed in relatively close proximity to adjacent negative voxels. There were also positively-weighted voxels in the right CST and in the splenium of the corpus callosum (Figure 4C). Given the directions of the relationships, manual dexterity was positively associated with the positive clusters in the left frontal, CST, and colossal WM while working memory was negatively associated with these regions (and vice versa).

We identified three clusters, two located in prefrontal WM (one positive and one negative) and one positive cluster in the CST. Across these clusters, FC, T1w/T2w (and in cluster 2, ICVF) contributed most strongly to the D2 scores (Figure 4D). The positive prefrontal cluster contained streamlines associated with the left superior medial frontal cortex, whereas the adjacent negative prefrontal cluster comprised streamlines connecting the left inferior frontal orbital cortex (Figure 4E). Finally, the positive CST cluster corresponded to the passage of streamlines connecting the right precuneus, thalamus, and parietal cortices.

**Figure 2:**
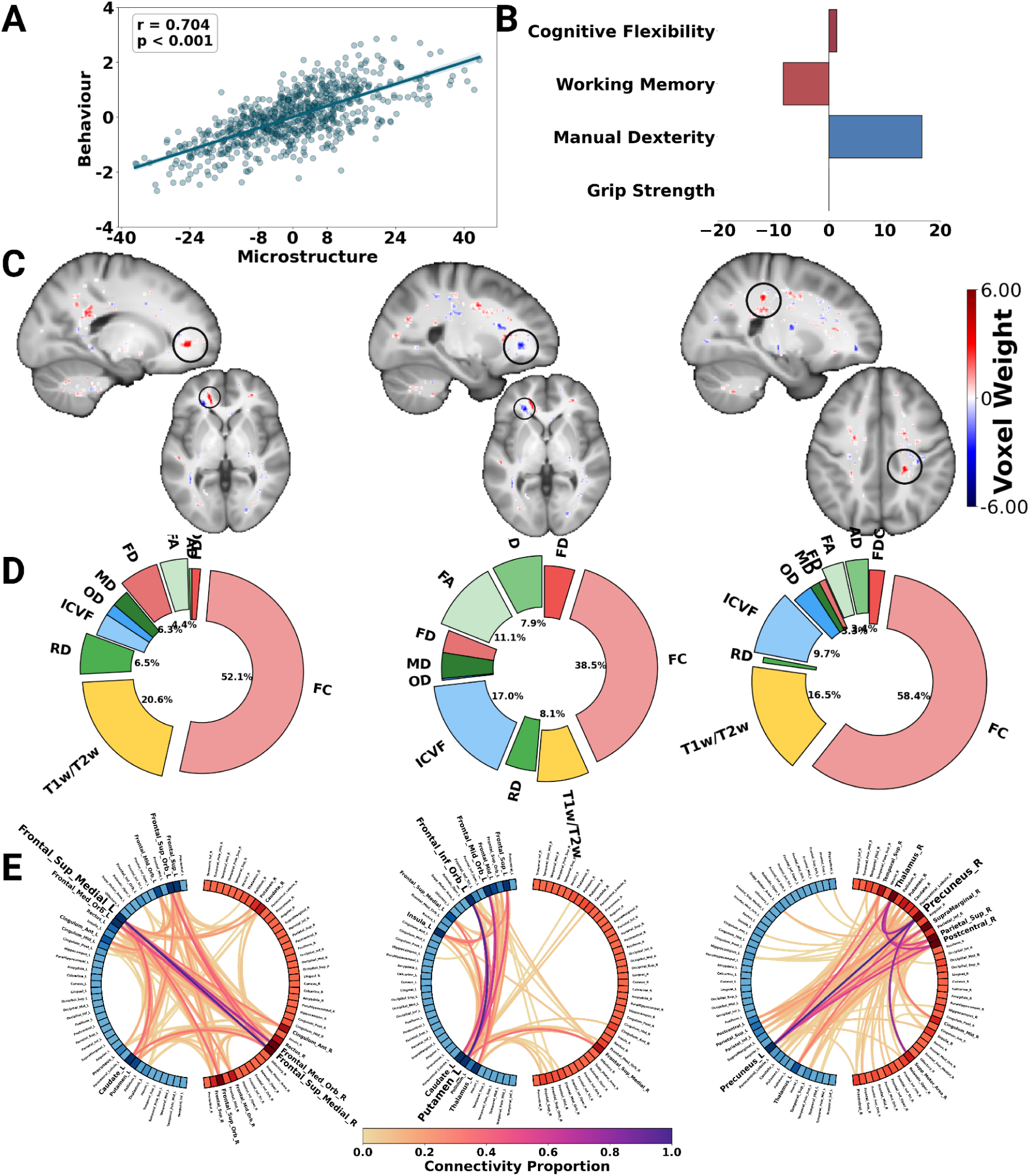
Decomposition results for the first latent variable. (A) PLS score correlation; (B) weights of statistically significant tasks; (C) weights of statistically significant voxels; (D) metric contributions to the D2 score; (E) connectivity profiles of spatially contiguous clusters based on the AAL atlas.

## 4. Discussion

The goal of the current study was to better understand how individual differences in white matter (WM) microstructure are related to differences in behaviour. We integrated 10 WM metrics into individual voxelwise distances from the population mean (D2) and then examined the spatial relationships between D2 and four behavioural tasks from two domains (cognition and motor). We used the MVComp toolbox (https://github.com/neuralabc/mvcomp) to compute voxelwise D2 scores from WM metrics while accounting for their covariance, and then extracted latent variables of the spatial pattern of association between D2 and behaviour using PLS-SVD. Our approach was designed to meet the need for a mapping of behaviour and WM microstructure using a holistic and integrative multivariate solution with high validity and reliability compared to traditional approaches (Varoquaux et al., 2018; Yoo et al., 2019). We identified three statistically significant latent variables (LVs) that explained a total of 83% of the brain-behaviour associations (Figure 1) that broadly decomposes this relationship into higher-order cognitive and premotor function (LV1; Figure 2), motor function (LV2; Figure 3), and integrative cognitive-motor function (LV3; Figure 4).

We identified statistically significant relationships between microstructure and behaviour in three of the four latent variables (panel a in Figure 2,3,4). Combining multiple cognitive and motor tasks allowed for the extraction of microstructural correlates of behaviour that take into account both the physiologically ambiguous and overlapping nature of imaging metrics (Jelescu & Budde, 2017; Novikov, 2021) and the complex mapping between behavioural constructs and task performance (Schöttner et al., 2023; Schuch et al., 2022; Yoo et al., 2019). Our findings identified three distinct patterns: a cognitive (LV1), a motor (LV2), and an integrative (LV3) behaviour pattern. This is in agreement with the current frameworks in cognitive neuroscience that focus on the complex modelling of behaviour, starting with (Price & Friston, 2005), and more recently expanded on in (Poldrack, 2010; Varoquaux & Poldrack, 2019). These authors suggested that past traditional methods of bivariate brain-behaviour assessments limit their interpretability and generalizability across behavioural domains. Our method avoids oversimplifying behaviour, and rather extracts latent variables that link performance on different behavioural tasks to individual differences in WM microstructure. Hence, our work provides a potential avenue for developing behavioural/task mappings beyond the cortex and into WM. Overall, the effects we observed could be due to neuroplastic changes in WM that lead to enhanced task performance. Although not causal, our technique shows that combining multiple measures of behaviour with multiple measures of microstructure provides a holistic and uniquely powerful window into how the brain supports behaviour.

Based on the behavioural weights of each latent variable, we interpret LV1 as representing higher-order cognitive and premotor skills, LV2 representing motor behaviour, and LV3 representing integrative functioning (panel B, Figure 2,3,4). Our results also emphasise the connectedness and interplay of different components of behaviour during a given task (Schöttner et al., 2023). For example, while the cognitive tasks (cognitive flexibility and working memory) load strongly on LV1, dexterity shows a similar weighting, suggesting a contribution from premotor planning. Supporting this interpretation, dexterity has been previously grouped with cognitive tasks via experimental manipulation (Rodríguez-Aranda et al., 2016) and when behavioural latent spaces are explored via machine learning (Schöttner et al., 2023). Similar to the pattern expressed by LV1, previous studies examining the neural correlates of fine motor skills suggest that many cortical regions are involved in this task, including visuospatial and somatosensory areas, such as premotor and primary motor areas (Sobinov & Bensmaia, 2021). In contrast, LV2 shows a trade-off between grip strength (strong positive) and dexterity (weak negative), with cognitive flexibility also weighing positively, while working memory, which does not require arm use, does not load significantly (Weintraub et al., 2013). In line with the task loadings on LV2, meta-analytical evidence from functional imaging during cognitive flexibility tasks shows activation in cognitive and motor areas of the cortex (Buchsbaum et al., 2005). This LV further separates gross motor (grip strength, cognitive flexibility) from fine motor (dexterity) movements (Figure 3B). Lastly, the third LV shows strong positive and negative weights of cognitive and motor tasks, reflecting its integrative nature. This integrative aspect could only be extracted with a multivariate approach that combines multiple behavioural domains. Thus the incorporation of WM microstructure in behavioural analysis provides a grounding for behaviour in neuroanatomy. While there is evidence for disrupted cognitive-motor integration in relation to cortical functioning in TBI (Sergio et al., 2020) and post-concussion (Hurtubise et al., 2020), to our knowledge this study is the first to decompose the multimodal relationship between healthy WM and tasks from different behavioural domains. Therefore, these results demonstrate that behavioural performance across multiple domains can provide meaningful insights into WM microstructure.

When examining the voxel weights, we observed widespread shared and overlapping patterns across WM in all LVs (panel C, Figure 2,3,4). These findings expand our understanding of the complexity of brain-behaviour associations, emphasising the active role of WM microstructure in supporting higher-order cognitive processes, motor skills, and the integration of cognitive and motor functions. This is evident in the voxels’ weights that span regions traditionally labelled as cognitive, motor, and higher-order integration. For instance, frontal and parietal WM for cognitive functions, parietal, cerebellar, and frontal WM for motor skills, and prefrontal WM for integrative functioning. First, the cognitive LV1 is mostly negatively distributed, indicating that better performance on cognitive tasks is associated with smaller deviations of WM microstructure relative to the reference population. This pattern is observed for anterior and posterior frontal, and parietal WM. Our results are in agreement with recent work on stroke-based disconnectomes, which shows impairments in both cognitive and dexterity skills with disconnection in these regions (Talozzi et al., 2023). Moreover, cortical regions involved in premotor functions, visuospatial imagery, and executive skills are connected through these WM regions (Krüger et al., 2020; Sobinov & Bensmaia, 2021; Suchan et al., 2002). Namely, the SMA, the precuneus, putamen, and frontal cortices, as shown in panel E in Figures 2, 3, and 4. Since our results come from a large sample of healthy individuals, we demonstrate that it is possible to use an individual difference approach in combination with multivariate decomposition to localise motor and cognitive function in WM in the absence of pathology. Detailed normative knowledge of this fundamental relationship between WM microstructure and behaviour could aid in symptom prediction and personalised treatment in diseases that affect WM such cardiovascular damage, Alzheimer’s Disease, and in stroke. Second, the motor LV (LV2) has positively- and negatively-weighted regions differentially across all of WM. The second LV’s positive voxels are highly associated with motor function, and are primarily distributed in WM underlying the left motor regions. We hypothesize that this is largely due to the recruitment of these regions in signal conduction, and the modulation of the microstructure for optimal motor performance (Sampaio-Baptista et al., 2013). The corticospinal tract was also positively associated with gross motor performance, and negatively with fine motor skills. In the current study, most tasks (with the exception of the working memory task) required motor execution, and the motor task that had the least cognitive requirements (grip strength, for which planning is not necessary) had the largest weight. This is also supported by the cortices most connected through these clusters. We observed maximal connectivity of the left thalamus, left and right peri-central and frontal superior cortices in these clusters. For the third latent variable (LV3), our results provide evidence for integration of motor function with cognitive abilities, including prefrontal WM, parietal WM, and the splenium of the corpus callosum. These findings are consistent with previous evidence linking motor planning (i.e., the dexterity task), executive function, and working memory functioning to prefrontal areas (Rodríguez-Aranda et al., 2016; Seidler et al., 2012; Thiebaut de Schotten & Forkel, 2022). Microstructural differences in these regions are likely to be part of a larger network of cortical regions connected through WM, since the extracted cortical regions indicate that prefrontal GM is involved in integrative functioning (Seidler et al., 2012). Therefore, our observations in the prefrontal WM may reflect a modulation of microstructure to support cortical functioning. Across all regions and the three identified LVs, these findings suggest that the observed behaviours may arise from a complex interplay between WM microstructure and connectivity, where structural properties shape information flow and functional integration across cortical networks.

This study is not without limitations. We used four tasks spanning cognitive and motor behaviours to provide a holistic brain-behaviour mapping; however, in order to provide a complete behavioural atlas grounded in WM microstructure, more tasks should be incorporated in the future. Schöttner et al. (2023) showed that four factors extracted from a large battery of tasks provided the optimal tradeoff between variance, reliability, and validity in behavioural assessments. It should be noted that adding more behavioral assessments may increase model complexity, potentially reducing interpretability of the results. Also, other functions, such as sensory and emotional behaviours, are not measured with these four tasks (Varoquaux & Poldrack, 2019). Therefore, future examinations should use similar dimensionality reduction techniques to Schöttner et al. (2023) while also incorporating emotional, language, and sensory measures. This will provide a more complete atlas of human behaviour, and disentangle the contribution of microstructural differences across WM in healthy function. Moreover, the Mahalanobis distance suffers from two main issues. First, it is a squared metric that does not distinguish the directionality of deviation from the average. Therefore, it is not possible to assess whether there has been an increase or a decrease relative to the reference population. Directionality could, for example, be an important factor in cases where it is of interest to assess longitudinal change, where the average is substituted as the baseline. However, it is possible to extract metric contributions to D2, enabling post-hoc assessment of directional differences. Second, voxelwise D2 values reflect how much a subject’s microstructure deviates from the reference average. While thresholds for significance have been proposed (Guerrero-Gonzalez et al., 2022; Penny, 1996), they rely on normality assumptions that are not always met. Future studies could also use permutation or reshuffling methods appropriate at the individual level to identify robust D2 deviations/longitudinal change to explore within subject microstructural changes and their relationships to behaviour.

## Conclusion

In conclusion, we provide a holistic mapping between cognitive and motor behaviours and white matter microstructure. We achieve this goal by using the Mahalanobis distance (D2) computed from 10 microstructural metrics, and decomposing the association between D2 and four behavioural measures assessing cognitive flexibility, working memory, dexterity and strength. We show that using multivariate statistical analysis reveals strong brain-behaviour associations and is able to extract latent variables of cognitive, motor and integrative function. We also provide spatial mappings between these three domains of behaviour and WM microstructure. We extend our analysis to the cortical GM, and show that there exists an interplay between connectivity and microstructure which gives rise to and sustains healthy human behavioural functioning.

## 5. Statements & Declarations

### Data and Code Availability

All data used in this study is available from the Human Connectome Project Database. The mvComp toolbox used to compute the voxelwise D2 maps is open-source and available at (https://github.com/neuralabc/mvcomp).

### Ethics approval & Consent to participate & Consent to publish

Data acquisition for this study was performed in line with the principles of the Declaration of Helsinki, and the HCP acquired informed consent prior to any data collection. All data used in this study was published with consent to publish from the HCP.

### Competing Interests

The authors have no relevant financial or non-financial interests to disclose.

### Author Contributions

Zaki Alasmar: Methodology, Software, Formal analysis, Validation, Conceptualization, Data Curation, Writing - Original Draft, Visualization. Stefanie A Tremblay: Writing - Review & Editing, Data Curation, Methodology, Software, Validation, Visualization. T. Robert Baumeister: Methodology, Software, Writing - Review & Editing. Amir Pirhadi: Methodology, Software, Validation, Data Curation. Felix Carbonell: Methodology, Writing - Review & Editing. Yasser Iturria-Medina: Methodology, Writing - Review & Editing, Conceptualization, Funding acquisition. Claudine J Gauthier: Supervision, Conceptualization, Writing - Review & Editing. Christopher J Steele: Supervision, Conceptualization, Methodology, Writing - Review & Editing, Software, Funding acquisition.

## Notes

### Competing Interest Statement

The authors have declared no competing interest.

### Summary of Updates

Updated language surrounding the results and their interpretation. Updated figure layouts and added methodological workflow figure.

